# Multi-Scale Neural Sources of EEG: Genuine, Equivalent, and Representative. A Tutorial Review

**DOI:** 10.1101/391318

**Authors:** Paul L. Nunez, Michael D. Nunez, Ramesh Srinivasan

## Abstract

A biophysical framework needed to interpret electrophysiological data recorded at multiple spatial scales of brain tissue is developed. Micro current sources at membrane surfaces produce local field potentials (LFP), electrocorticography (ECoG), and electroencephalography (EEG). We categorize multi-scale sources as *genuine*, *equivalent*, or *representative*. *Genuine sources* occur at the micro scale of cell surfaces. *Equivalent sources* provide identical experimental outcomes over a range of scales and applications. In contrast, each *representative source distribution* is just one of many possible source distributions that yield similar experimental outcomes. Macro sources (“dipoles”) may be defined at the macrocolumn (mm) scale and depend on several features of the micro sources—magnitudes, micro synchrony within columns, and distribution through the cortical depths. These micro source properties are determined by brain dynamics and the columnar structure of cortical tissue. The number of *representative sources* underlying EEG data depends on the spatial scale of neural tissue under study. EEG inverse solutions (e.g. dipole localization) and high resolution estimates (e.g. Laplacian, dura imaging) have both strengths and limitations that depend on experimental conditions. The proposed theoretical framework informs studies of EEG *source localization*, *source characterization*, and *low passes filtering*. It also facilitates interpretations of brain dynamics and cognition, including measures of *synchrony*, *functional connections between cortical locations*, and other aspects of *brain complexity*.

## 1. Introduction

### 1.1 Varying interpretations of brain “sources”

This review of EEG source biophysics is partly motivated by a comprehensive report, *International Federation of Clinical Neurophysiology (IFCN) guidelines for topographic and frequency analysis of resting state electroencephalographic rhythms* (Babiloni, 2018), produced by a Working Group of 15 EEG scientists (including author PLN). The report deals with various measures of neocortical dynamics, including source synchronization, functional connectivity, and various other brain complexity measures. Relationships between recorded EEG data and the underlying “sources” were considered, and the discussion revealed a number of controversies, to be expected when complex topics are considered. In particular, some disagreements apparently originated from divergent and/or poorly defined ideas of just what is actually meant by the tag “EEG sources,” which is the main topic of this paper.

Electric potentials are generated by brain sources; the central goal is to gain information about sources in relation to cognitive or clinical states. Such information might involve resting state EEG or event related potentials (ERP). Common measures include synchrony, functional connectivity, and various aspects of brain complexity in extended networks of sources. Despite wide recognition of the importance of brain source estimates, relationships between recorded data and the underlying sources are often obscure. Basic biophysical studies emphasize that brain “sources” should be defined at scales that suitably match the chosen measurement scales (Nunez, 1981, 1995; Srinivasan, 1999; Nunez and Srinivasan, 2006, 2014; Nunez, 2012; Sporns, 2011). The practice of electrophysiology requires many additional considerations, including reference electrode, head model, electrode density, noise, and artifact. But, here we mostly avoid these important topics in order to focus on a fundamental biophysical source framework. While much future progress in electrophysiology, including new computational and statistical developments, can be expected, the fundamentals of multi-scale brain sources presented here should remain largely unchanged for the foreseeable future.

### 1.2 Non uniqueness of source solutions

Many studies of EEG “source” localization have been published, even though such inverse solutions are well-known to be non-unique and thereby subject to a range of interpretations (Nunez, 1981, Nunez and Srinivasan, 2006). This background raises basic questions of how to evaluate such source representations. In order to shed more light on this and related issues, we propose distinctions between *genuine sources*, *equivalent sources*, and *representative sources*. In our chosen terminology, *genuine sources* consist of the current distributions at the micro scales of small parts of cell surfaces. *Equivalent sources* provide identical outcomes in experiments carried out over a range of scales and applications. In contrast, each *representative source distribution* provides just one of many possible configurations that could generate similar data at large scales, but might represent little more than a hypothesis. These source distinctions inform additional effects, including low pass filtering in ECoG versus EEG and different kinds of brain correlations, measures of functional connectivity like coherence and covariance between cortical locations.

### 1.3 Multi-scale source estimates

Brain electric potentials are recorded over a broad range of spatial scales determined mostly by the size and location of the recording electrodes. This occurs because the experimental data reflect potentials space-averaged over tissue volumes equal to or larger than the volume of the electrode tip. In the case of scalp potentials, the tissue-averaging volume is much larger than the electrode volume, because of the scalp electrode’s large distance from sources and the smearing effect of the intervening tissue, especially the skull. Potentials recorded from the cortical surface with ECoG arrays are also space-averaged, but over much smaller volumes than scalp potentials. Thus, one may define four distinct recording scales (Nunez, 1995, 2012; Nunez and Srinivasan, 2006):

- *Individual neurons*. *Micro scale* recordings of surface or trans-membrane potentials.
- *Local field potentials* (LFPs). *Small scale fields* recorded within brain tissue (usually cortical), mostly reflecting current sources due to synaptic activity occurring within perhaps 0.1 to 1 mm of the recording electrodes; that is, within tissue volumes typically in the 10^-3^ to 1 mm^3^ range.
- *Intermediate* (*meso*) *scale fields*. The *electrocorticogram* (ECoG) is recorded from the cortical, pia, arachnoid, or dura surfaces. Depending on specific location and electrode size, these potentials appear to reflect synaptic and other source activity occurring over some portion of the depth of local cortex (2 to 5 mm); that is, within tissue volumes of perhaps 1-20 mm^3^. In addition, epileptic patients may be implanted with *intracranial depth* (iEEG) electrodes to establish or refute the occurrence of seizure foci.
- *Macro scale fields.* Potentials recorded by the electroencephalogram (EEG) are obtained from the scalp; each electrode reflects synaptic source activity occurring within large parts of the underlying brain, something like 10 to 50 cm^2^ of the cortical sheet or cortical tissue volumes approximately in the 10^3^ to 10^4^ mm^3^ range (Nunez and Srinivasan, 2006). Thus, EEG typically represents the space-averaged source activity in tissue containing on the order of 100 million to a billion neurons.

Synaptic and action potentials at neural membranes create micro current sources, the so-called *generators* of LFP, ECoG, and EEG signals. These same current sources also generate magnetic fields (MEG), which have different sensitivity to specific source characteristics (Hamaleinen et al, 1993; Srinivasan et al, 2007). At the low frequencies that are of interest in electrophysiology, the electric and magnetic fields are *uncoupled*; that is, each may be estimated without reference to the other (Nunez and Srinivasan, 2006, appendix B). For this reason, we avoid the label *electromagnetic* which implies a single (coupled) field that generally exhibits more complicated dynamic behaviors.

While large-scale measures like EEG inform the big picture but provide almost no local details, small-scale measures like LFPs provide local detail but only sparse spatial coverage. Much of the membrane current from source regions remains in the local tissue and forms small, closed current loops that pass through the intracellular, membrane, and extracellular media. Such local source activity may be recorded as LFP. In addition, some of the same source current may reach the cortical surface to be recorded as ECoG, and a little even gets as far as the scalp to be recorded as EEG. The manner in which source current spreads through brain, CSF, skull, and scalp tissue is labeled *volume conduction*; it is determined by the geometry (tissue surface boundaries) and electrical *conductivity* (or its inverse, *resistivity*) of these tissues. Thus, these measures, plus the intermediate-scale ECoG, provide *complementary* and largely independent measures of brain source activity at different spatial scales, and therefore must employ different levels of description. This independence arises partly because larger scale measures are selectively sensitive to the smaller scale *synchronous* source populations, whereas the *asynchronous* micro source potentials tend to cancel in larger scale measurements (Nunez and Srinivasan, 2006).

## 2. Equivalent sources in physical networks

Before exploring distinct categories of brain sources, we introduce similar ideas from physical networks in electrical engineering applications. We show that, similar to brain sources, physical voltage or current sources may be considered *genuine*, *equivalent*, or *representative*. Figure 1a depicts a large network separated into two sub-networks **L** and **E** connected at the two ports a and b. Each network might consist of thousands of circuit elements— current sources, voltage sources, resistors, capacitors, and so forth. The sources may be independent or dependent on voltages or currents at other locations in the same sub-network. For our purposes, voltage measurements within networks **L** and **E** are analogous to LFP and EEG recordings, respectively, and are reflected by the cartoon head image. A technical aside—we assume the **L** network to be approximately linear to simplify the proposed analogy, whereas the **E** network may be nonlinear.

**Figure 1.**
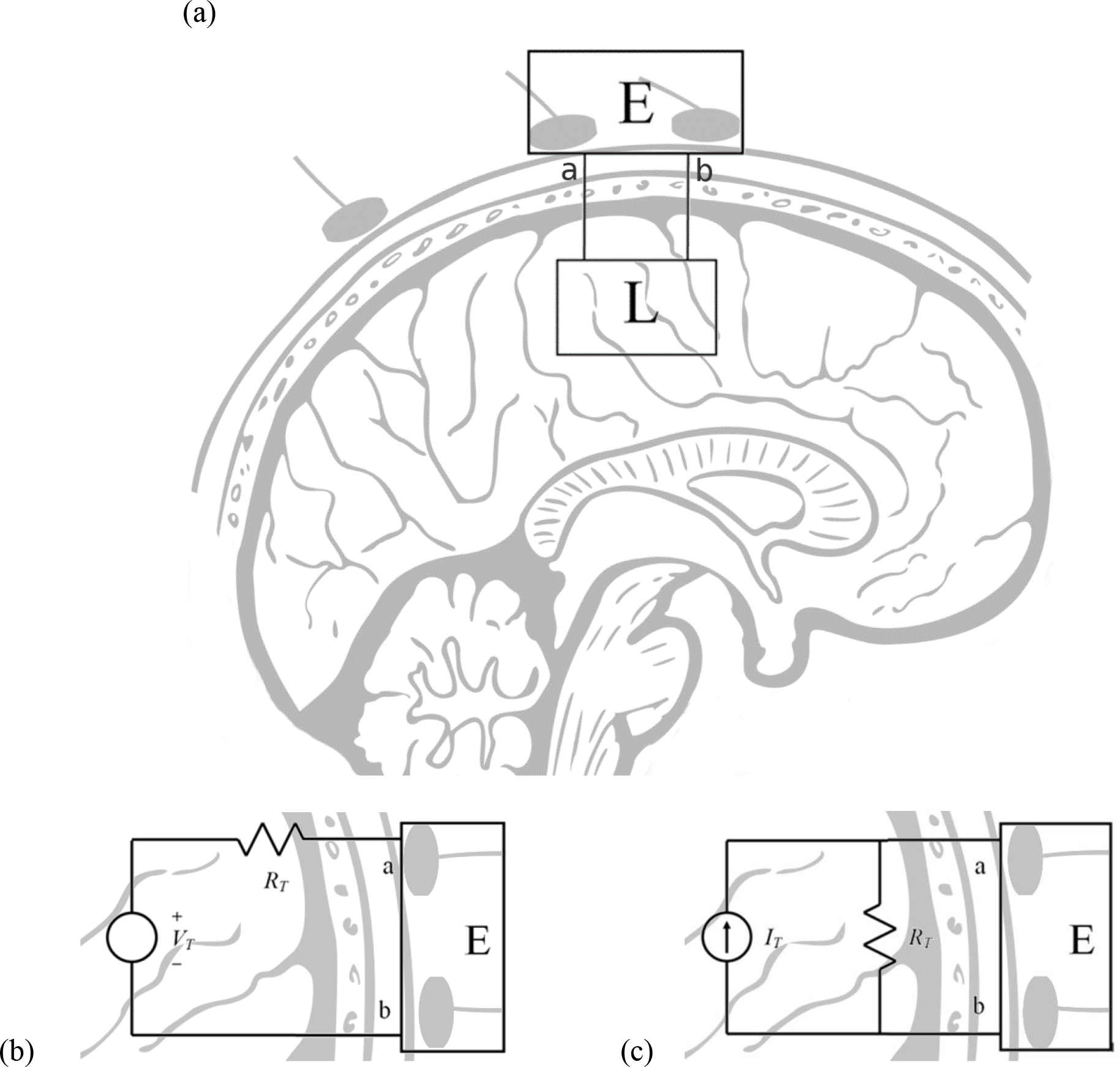
(a) Two electrical sub networks **L** and **E** are connected at the ports a and b. The background brain cartoon emphasizes the analogues, but does not influence the circuit discussions. (b) Thevenin equivalent network. (c) Norton equivalent network. Networks b and c are “equivalent” to each other and to network a in the sense of producing exactly the same currents and voltages at all locations in network **E**.

Figure 1b contains an ideal independent voltage source (battery or AC generator *V*_*T*_), and fig.1c contains an ideal independent current source *I*_*T*_. “Ideal independent” means that the magnitude of the voltage or current produced by each independent source is a fixed property and is not affected by other elements in the circuit. Here the symbol *R*_*T*_ indicates impedance (for AC circuits) or just its real part (resistance) since the imaginary part of impedance (due to capacitive effects) is typically negligible in large tissue volumes; however, none of our findings depend on this finding. Thevenin’s theorem of electrical engineering says that the network of fig. 1b, consisting of a voltage source *V*_*T*_ in series with a resistor *R*_*T*_, is “equivalent” to network **L** (fig. 1a) in the following sense: All the currents and voltages in network **E** are unchanged, regardless of whether the terminals (*a*, *b*) are connected to network **L** or to the simple voltage source and series resistor of fig. 2b. Similarly, network **L** may be replaced by the current source *I*_*T*_ and parallel resistor *R*_*T*_ shown in fig. 2c, where the equivalent voltage source and equivalent current sources are simply related by *V*_*T*_ = *R*_*T*_ *I*_*T*_. The networks of figs. 2b and 2c are called the *Thevenin and Norton equivalent networks*, respectively (Nilsson, 1986, or nearly any other book on electric circuits).

**Figure 2.**
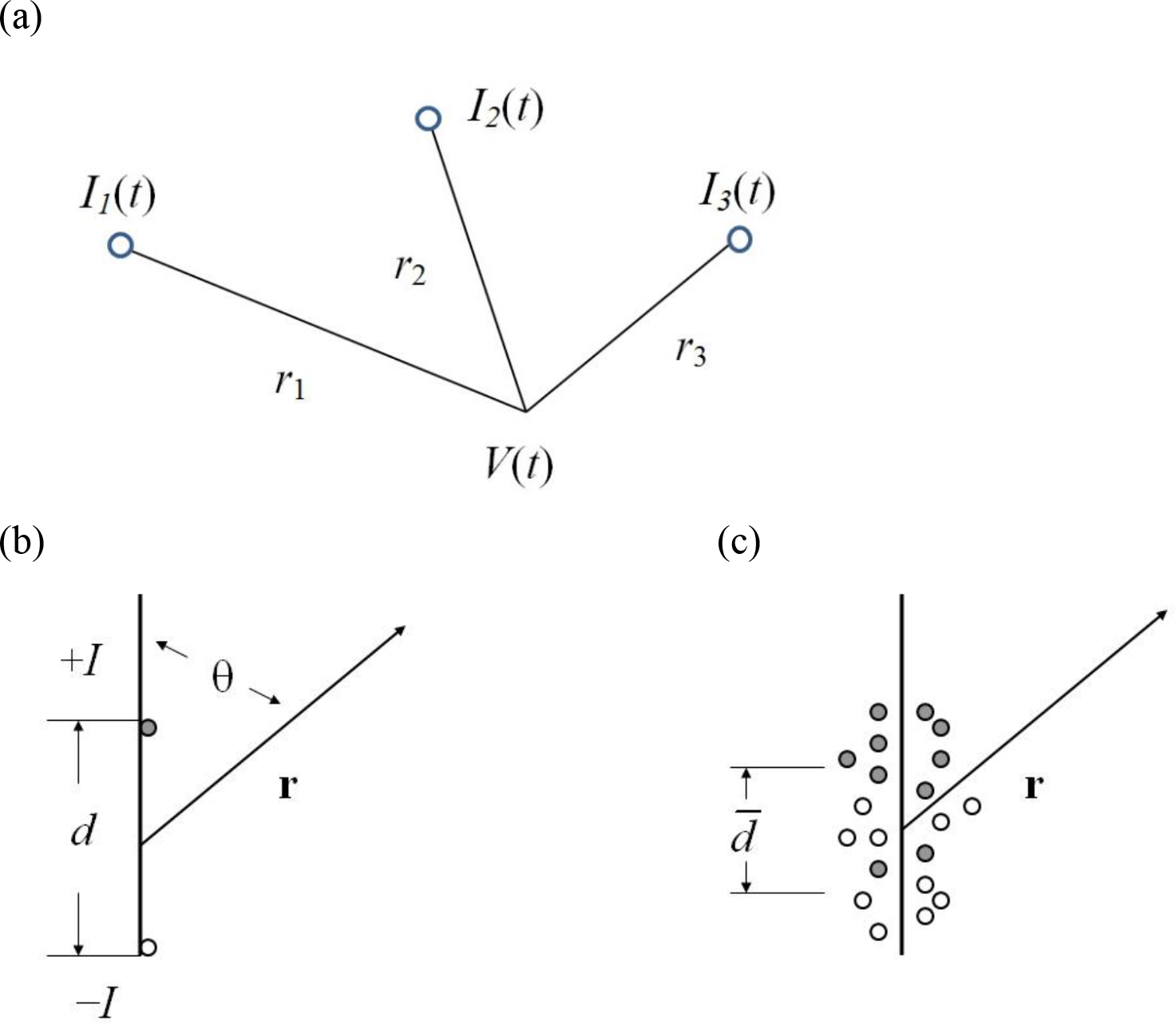
Monopole and dipole source distributions and the resulting potentials *V*(*t*). The filled and empty circles represent positive and negative point current sources, respectively. (a) isolated monopolar sources (b) simple dipole (c) distributed point sources along a vertical axis.

If we are only interested in what happens inside network **E**, we can accurately replace network **L** by its Thevenin or Norton equivalents shown in figs. 2b and 2c. However, the idea of source “equivalence” can be carried too far. For example, what does perfect knowledge of the Norton equivalent source of fig. 2c tell us about the actual internal voltages within network **L**? The answer is almost nothing except that one or more sources at unknown locations must be active within **L**. This inverse problem is non-unique—an infinite number of **L** networks, possibly containing millions of independent and dependent sources, will produce the identical currents and voltages in **E**. If we are interested in what happens at various locations within network **L**, this network must be studied in its original form. Analogous arguments may be applied in brain tissue. For example, suppose we find the magnitude of a “macro source” in a cortical column (defined in section 4). Each brain macro source consists of billions of synaptic scale micro sources acting at membrane surfaces (analogous to the sources in **L**). As in the case of the physical networks of fig. 1, the macro sources can be labeled *representative* or for some limited purposes *equivalent*, but not *genuine*, because even perfect knowledge of the macro sources tells us little about the underlying micro source details.

## 3. Micro sources at membrane surfaces

### 3.1 Monopole sources

Any current source region in a volume conductor like brain tissue may be represented (modeled) as a sum of distributed point sources and sinks, as indicated in fig. 2a. The potential due to *N* point (monopolar) sources *I*_*n*_(*t*) in an idealized homogeneous and isotropic medium of conductivity σ is given by (Nunez and Srinivasan, 2006; Nunez, 2012),

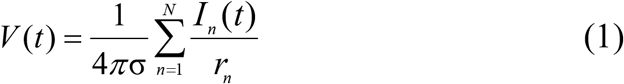

Equation (1) follows directly from law of current conservation and Ohm’s “law.” Current conservation is a fundamental law; Ohm’s law not really a law, but is expected to provide good approximations to macro scale tissue volumes (Nunez and Srinivasan, 2006). A numerical example for brain tissue is thus—let a single current source *I*_n_ be 4*π* microamperes (μA), and let cortical resistivity (inverse of conductivity σ) be 3,000 ohm mm. The predicted potential in the cortex on a spherical surface of *r* = 1 mm radius surrounding a single point source is then 3,000 μV, assuming all other sources and sinks are located much farther away (but note the caveats below). In genuine neural tissue, however, such sources may be partly or mostly cancelled by nearby sinks resulting in much lower potentials. Equation (1) incorporates several idealizations and requires the following caveats:

- *V*(*t*) indicates the potential with respect to infinity, approximated with a “distant” (compared to the source region) reference electrode.
- All current sources must be balanced by current sinks somewhere in the volume conductor as required by current conservation.
- There is a distinction between the point potential of Eq (1) and the potential recorded with a real electrode of non-zero radius; that is, measured potentials represent space-averages over the electrode tip volume.
- The medium is assumed to be infinite with constant scalar conductivity σ; that is, no boundary or direction-dependent tissue effects are present.

In one example, the monopolar representation of Eq (1) may be employed to estimate the extracellular potential fall-off of the action potential. This extracellular potential may be *represented* by sub-millimeter monopolar source and sink regions using Eq (1), approximately replicating the triphasic waveform of the action potential and forcing total membrane source current to equal total sink current (Nunez and Srinivasan, 2006, chapter 5). Distributed monopolar rather than dipolar source models are required because of the large (cm scale) source-sink separations of the triphasic action potential in myelinated axons. The resulting model predictions provide a reasonable match to an experiment with the compound action potential of the frog sciatic nerve (Flick et al, 1977). Such *representative* source distributions can be expected qualify as *equivalent* for some limited experimental measurements at macroscopic (cm) scales. However, genuine studies of action potential generation require smaller scale and more detailed studies of nonlinear membrane properties (Cole 1968).

### 3.2 Dipole sources

When many point sources are present, application of Eq (1) can be quite cumbersome, as in the above example of action potential sources. However, in other cases, all sources and sinks may be confined to a region that is much smaller than the nearest distance to measurement points. If so, the distant potential generated by the source-sink region may be approximated by a dipole expression that is much simpler to use than Eq (1). The idealized *current dipole* consists of a point source *+I* and a point sink −*I*, separated by a distance *d*, as shown in fig. 2b. However, the word *dipole* has a more general and useful meaning, making the dipole concept applicable to a wide range of source-sink configurations. Nearly any source-sink region where the total source and sink currents are equal (local current conservation) will generate a predominantly dipole potential at distances that are large compared to the dimensions of the source-sink region. Thus, the collection of point sources and sinks shown in fig. 2c produces an approximate dipole potential at distances *r* when *r* is large compared to *d*, in practice greater than perhaps 3*d* or 4*d* depending on desired accuracy. For this reason, cortical dipoles, and especially dipole layers (sheets) of various sizes, provide useful source models for potentials recorded on the scalp. If all the sources occur near the vertical axis, the potential due to either of the source distributions in figs. 2b or 2c may be approximated by (Nunez and Srinivasan, 2006; Nunez, 2012),

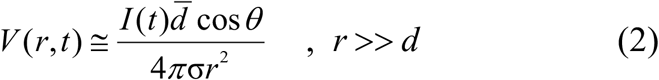

Equation (2) involves only a single distance *r* between the measuring point and the center of the source-sink region, rather than Eq (1), which might involve millions of distances *r*_*n*_. Here *θ* is the angle between the (vertical) dipole axis and the vector **r** to the point of measurement. The “effective pole separations” are given by the symbol *d̅*, which accounts for the mixing of positive and negative point sources. In the case of the simple dipole of fig. 2b, *d̅* = *d*; whereas *d̅* < *d* for the distributed sources in fig. 2c. For example, if a point source, perhaps simulating an inhibitory postsynaptic potential (IPSP) at a cell body, is combined with passive sinks distributed uniformly over a distance *d* (the passive return current), the effective pole separation of the dipole is *d̅* = *d*/2 so the generated potential at large distances is exactly half as large as in the simple dipole of fig. 2b. The angular dependence in Eq (2) is strictly correct only if all sources lie on the vertical axis. However, Eq (2) provides a reasonable approximation if the sources approximate a narrow cylindrical region as shown on the right side of fig. 2c, perhaps a small diameter cortical column.

### 3.3 Open and closed fields

As indicated above, when the average separation between point sources and sinks is reduced, the effective pole separation and external potential also become smaller. The so-called *closed field* of electrophysiology corresponds to the limiting case *d̅* → 0, which occurs when positive and negative point sources are well-mixed, having small average separations. Interestingly, this means that the layered structure and synaptic distribution within mammalian cortex is critical to the production of scalp potentials—little or no recordable EEG would be expected if cortical neurons were randomly oriented or if excitatory and inhibitory synapses were fully mixed through cortical columns. Similarly, dipole magnitudes tend to be small if sources are asynchronous within the source region. In other words, excessive source activity within some local tissue volume need not produce a large dipole source if the underlying micro sources are mixed or asynchronous within the tissue voxel. This feature has implications for EEG/fMRI co-registration studies, suggesting that large fMRI signals need not “match” large EEG sources in the same tissue volumes (Nunez and Silberstein, 2000).

## 4. Macro source models

### 4.1 Macro sources in cortical columns

In classical electromagnetic theory, the importance of matching theoretical and experimental scales is widely appreciated (Jackson, 1975; Nunez, 1981, 1995; Nunez and Srinivasan, 2006). For example, different micro and macro electric and magnetic fields are defined and appear in the distinct micro and macro versions of Maxwell’s equations, which govern the behaviors of electric and magnetic fields in all material media, including tissue. To cite one common case— when macroscopic charges are placed in a dielectric (insulating material), numerous atomic scale charges rearrange themselves and create a complicated microscopic electric field pattern that strongly alters the macroscopic electric field. This example of micro scale charges in dielectrics is closely (in a mathematical sense) analogous to neural micro current sources in tissue, although the underlying physics is quite different. Macro scale tissue volumes behave mostly like conductors, but they also exhibit important dielectric properties. While some neuroscientists appear to view the issue of spatial scale as a minor inconvenience, we suggest it is critical to many applications involving EEG and the interpretations of such data (Nunez, 1981, 1995; Nunez and Srinivasan, 2006, 2014).

Each active synapse produces local active membrane current plus passive return current from more distant membrane surfaces as required by current conservation. Excitatory synapses produce local sink regions and distributed positive sources at more distant membrane locations. Inhibitory synapses produce current in opposite directions, that is, local membrane sources and more distant distributed sinks. Given the extreme complexity of the billions of micro scale sources in each macro scale tissue volume, it is convenient to define a macro scale source measure to more conveniently characterize EEG sources. Regardless of type of source, we may express source current per unit volume *s*(**r**, **w**, *t*) produced in a small (synaptic scale) tissue volume Δ*W* within the macro volume *W* by

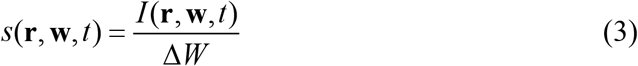

The vector coordinate **w** locates each synaptic scale current source (including passive return current) within a macroscopic tissue volume *W*, which may or may not be a cortical column. The vector coordinate **r** locates the center of the macro volume within the cortex, as shown in fig. 3. **P**(**r**, *t*) is defined as the *current dipole moment per unit volume*, given by the following triple integral over the tissue volume *W*

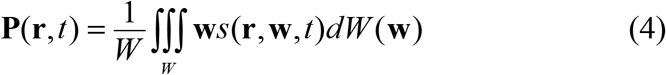

**Figure 3.**
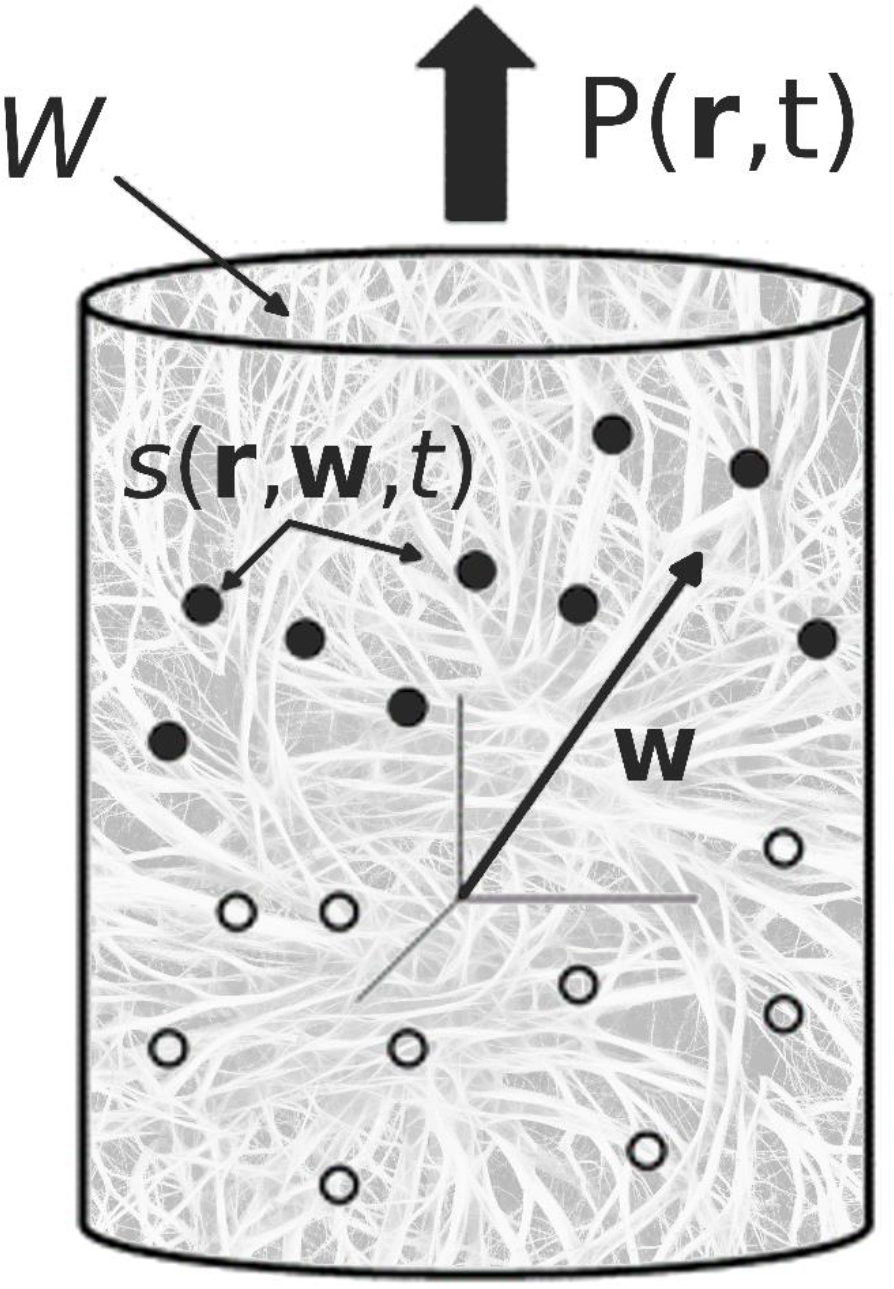
Micro sources *s*(**r**, **w**, *t*) in a neural mass *W* (e.g. a cortical column) produce macro sources **P**(**r**, *t*). The vectors **w** and **r** locate the micro sources and mass W, respectively.

### 4.2 Local current conservation?

This mathematical formalism applied to conductive media is identical to that of *charge dipole moment per unit volume* defined in dielectrics (insulators) (Jackson, 1975; Nunez and Srinivasan, 2006), although the physical basis is quite different (Technical point—both cases involve solutions to Poisson’s equation, which follows from Maxwell’s equations). The integral in Eq (4) may be applied to brain tissue volumes *W* of any size, including minicolumns, macrocolumns, or even the entire brain. However, for **P**(**r**, *t*) to be useful in the sense of providing genuine connections to EEG, several conditions must be met. First, the tissue volume *W* should be large enough to contain many micro sources *s*(**r**, **w**, *t*) due to local synaptic activity as well as the passive return currents. If, for example, the micro sources are defined at the scale of individual synapses, each (mm scale) macrocolumn would contain something like 10^10^ micro sources—a million neurons, each with ten thousand synapses plus an equal number of synaptic-sized cell patches for return current. This condition suggests that the total strength of micro sources may be approximately balanced locally by an equal strength of micro sinks such that the monopole contribution of the tissue volume *W* is approximately zero, that is

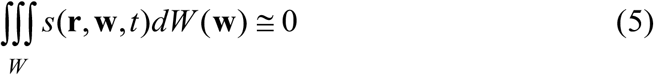

Comparison of Eq (5) with Eq (4) shows that the macro source strength **P**(**r**, *t*) depends on the particular manner in which the micro sources are distributed through the depth of cortex. In equivalent mathematical terms, the micro source function in the integral in Eq (4) is weighted by the location vector **w**. Thus, Eq (4) is just a generalization of the simple dipole expression, Eq (2). If the micro sources and sinks are randomly mixed or asynchronous within volume *W*, **P**(**r**, *t*) will be small or zero, even when the micro sources are large and numerous. While the local current conservation condition indicated by Eq (5) seems quite plausible, there is no guarantee that it is fully accurate in local tissue volumes. *Should this condition be inaccurate, the whole idea of macro scale cortical dipole sources would be called into question*. If, for example, action potentials in white matter (myelinated) axons, which may span cm scales, were to contribute substantially to EEG, the dipole model of Eq (4) would fail as a predictor of scalp potentials.

### 4.3 Implications for EEG versus ECoG

For EEG purposes, a second restriction on the chosen volume size *W* is this—all internal source-sink separations should be much smaller than the distances between the volumes *W* and the scalp. The maximum separation between upper and lower sources within a cortical column is about 5 mm, and the shortest distance between the center of a column and the scalp is perhaps 1.5-2 cm. In the dipole approximation of Eq (2), these estimates yield *r*/*d* = 3-4, suggesting that the dipole approximation is justified for rough approximations of cortical sources of EEG. With these restrictions in place, scalp surface potentials may be calculated based on the genuine dipole macro source **P**(**r**, *t*), with monopole, quadrupole, and higher ordered pole contributions negligible at the scalp (Jackson, 1975; Nunez and Srinivasan, 2006; Nunez, 2010a, 2012). However, in the case of ECoG, the electrodes are much too close to the cortical tissue *W* for the dipole approximation to be valid. Thus, any dipole source model, estimated only with EEG, provides severely distorted information about the sources associated with ECoG, which could be dominated by local monopolar sources in upper cortical layers. Perhaps more importantly, any local **P**(**r**, *t*) estimate based on scalp data will be influenced by nearby (or even distant) cortical activity. *Thus, we conclude that EEG and ECoG are not expected to be simply related; they provide distinct and complementary levels of description. This result emphasizes that the various common experimental measures of the brain’s dynamic behavior, including source localization and functional connectivity cannot generally be viewed in absolute terms; such measures are expected to be scale-dependent*.

### 4.4 Selective sensitivity of recording methods

For the reasons listed above, Eq (4) appears to be most useful in forward and inverse modeling of EEG when the chosen volume size *W* lies roughly between the minicolumn and macrocolumn scales. In either case, distances between cortical micro sources are less than 5 mm so the dipole approximation holds approximately. If the macrocolumn scale is chosen in EEG applications, the 1014 or so synaptic sources in all of neocortex may be represented by about ten thousand to a hundred thousand or so cortical “dipoles” (macro sources) forming a large dipole sheet, spread over the entire cortex (in and out of cortical folds). Of course, many regions may make negligible contributions to scalp potentials, depending on brain state, distances between sources and electrodes, and cancelling effects like macro sources active on opposite sides of cortical folds. In any case, neocortex may be treated as a continuum so that the macro source function **P**(**r**, *t*) is a continuous *field variable* forming a folded dipole sheet.

Figure 4 depicts neocortical sources forming *dipole layers* (dipole sheets) in and out of cortical fissures and sulci. Here each arrow indicates a macrocolumn scale source, a discrete representation of the function **P**(**r**, *t*) with strength varying as a function of cortical location. EEG is most sensitive to the correlated dipole layer in gyri (regions a-b, d-e, g-h), less sensitive to the correlated dipole layer in one sulcus (region h-i), and relatively insensitive to the opposing dipole layer in sulci (regions b-c-d, e-f-g) and the random layer (region i-j-k-l-m). ECoG is expected to record mostly superficial sources in local gyri. MEG is most sensitive to the correlated and minimally apposed dipole layer (h-i) and less sensitive to all other sources shown, which are opposing, random, or radial dipoles (Nunez, 1995; Srinivasan et al, 2007). In section 8, we argue that source selectivity also applies to high resolution EEG (HR EEG), which records at an intermediate scale between ECoG and EEG. In summary, *EEG, HR EEG, ECoG, and MEG are all selectively sensitive to different sets of cortical source distributions. There is no a priori reason to expect them to closely match each other*, *although substantial overlap may occur in many cases.*

**Figure 4.**
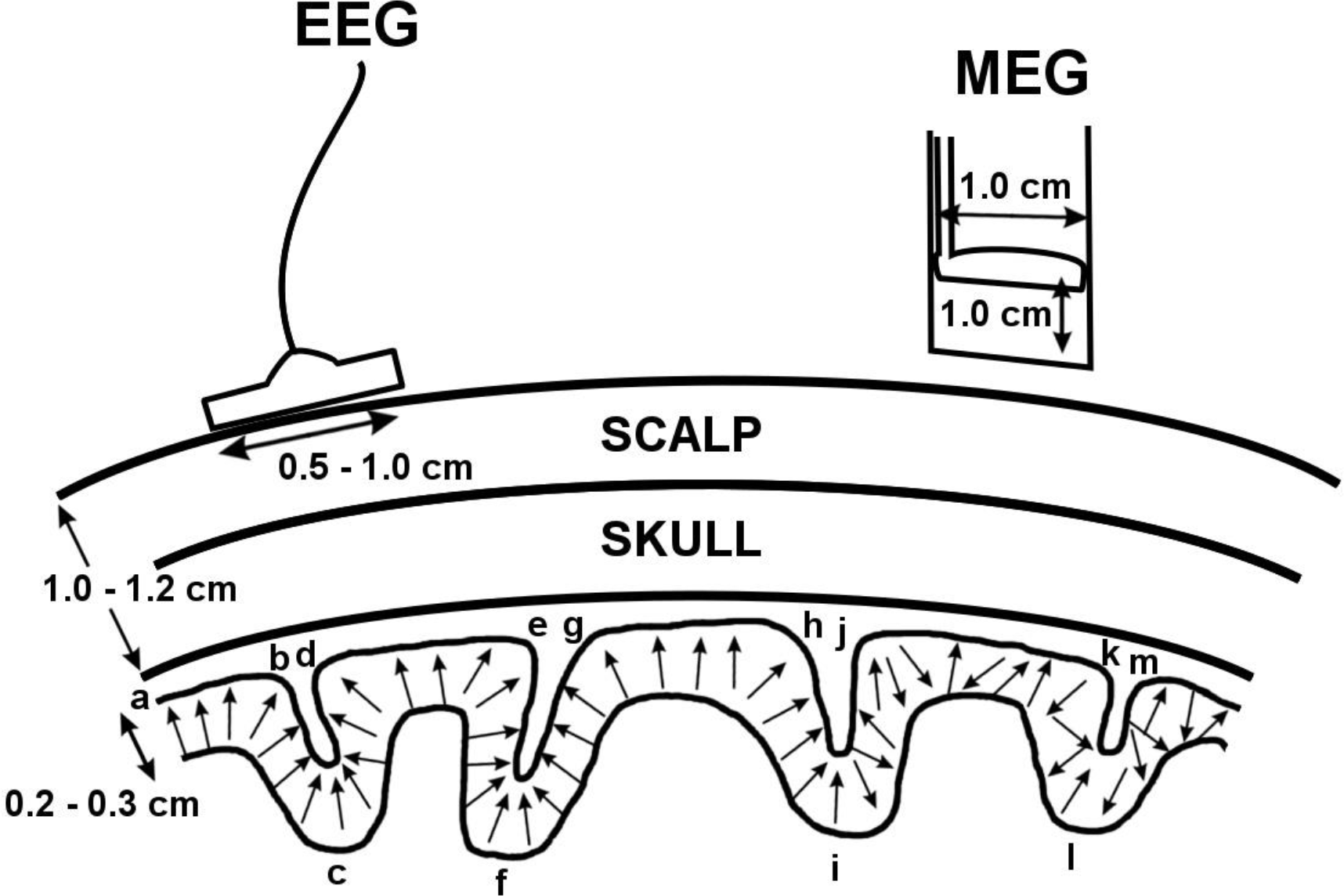
Cortical dipole layers. The arrows represent a snapshot of the macro source function **P**(**r**, *t*), which is here assumed to be synchronous and directed perpendicular to the local cortical surface over the extended region a-i. In contrast, **P**(**r**, *t*) has random directions in regions i-m.

The macro source function **P**(**r**, *t*) has the units of current density—micro amps per square mm (*μA*/*mm*^2^). In idealized cases, for example a macrocolumn of diameter 2 mm with all the sources in the lower cortex and all the sinks in the superficial cortex, **P**(**r**, *t*) is essentially the diffuse current density across the cortex (Nunez and Srinivasan, 2006). More generally, sources and sinks are expected to be mixed across the depths of the macrocolumn. In summary, according to Eq (4), the magnitude of **P**(**r**, *t*) depends on: (1) the sizes of the source-sink separations as in the simple dipole of Eq (2). (2) the micro synchrony of synaptic or action potential sources *s*(**r**, **w**, *t*) across the depth of columns—how closely they turn on and off together. (3) the numbers and magnitudes of the sources *s*(**r**, **w**, *t*).

## 5. Low Pass Filtering in ECoG and EEG

### 5.1 Source-based filtering of high frequencies

It has long been appreciated that scalp recorded EEG amplitudes tend to fall off at frequencies above the upper alpha band (13 Hz); with the higher frequencies sometimes dismissed as “1/*f* noise.” On the other hand, more recent ECoG recordings from human cortex have revealed a wealth of information about the possible functional roles of cross-frequency coupling involving frequencies up to perhaps 150 Hz (Canolty et al, 2010; Canolty and Knight, 2010). Some implications of these results have been discussed employing mathematical models (Nunez, 1974, 1989, 2010b, 2016; Nunez and Srinivasan, 2010, 2014; Srinivasan et al, 2013). The differences between EEG and ECoG magnitudes at high frequencies are partly (or perhaps mainly) due to the fact that higher frequencies tend to be less synchronous over the cortex than lower frequencies. As a result, the volume conductor (especially skull and scalp) acts as an “electroencephalographic averager.” Essentially, the temporal filtering observed at the scalp is a byproduct of spatial filtering caused by cortical dynamic behavior (Cooper et al, 1965; DeLucchi, 1975; Pfurtscheller and Cooper, 1975; Nunez, 1989, 1995; Nunez and Srinivasan, 2006).

An additional filtering mechanism, separate from this tangential synchrony over the cortex, may act within cortical columns, one which may also contribute to low pass temporal filtering at the scalp. Based on the classical cable theory of axons (Cole 1968), typical source-sink separations depend on the capacitive-resistive properties of cell membranes, predicting smaller source-sink separations (*d̅* small) at higher field frequencies. This implies a low pass frequency effect, reducing **P**(**r**, *t*) and as a result, lower scalp recorded EEG amplitudes at high frequencies (Nunez and Srinivasan, 2006, chapter 4). On the other hand, this low pass effect on scalp EEG magnitudes (due to reduced source-sink separations) might be much less important in ECoG recordings if ECoG potentials are mainly due to nearby monopolar sources in superficial cortex, modeled by Eq (1). This membrane scale capacitive-resistive influence on source-sink separations is quite different from the capacitive effects that might occur in large scale tissue volumes. The former (columnar) effect predicts amplitude reductions of EEG potentials at progressively higher frequencies caused by reduction in source-sink separations across the cortex. Simply put, the typical effective source separation *d̅* in Eq (2) or its generalization in Eq (4) is reduced because of cell membrane behavior at high frequencies, thereby lowering **P**(**r**, *t*). By contrast, bulk tissue capacitive affects result only in phase shifts (typically very small) between sources and potentials given by a non-negligible imaginary part of the tissue impedance.

### 5.2 Implanted artificial dipoles

To make this important distinction between separate capacitive influences more clear, consider the following experiment: An artificial dipole was implanted in cortex producing AC source current, representing **P**(**r**, *t*) (Cooper et al, 1965). The resulting magnitude of potential on the scalp was measured as a function of source frequency. If any measureable bulk capacitive effects were to occur, they would be observed as phase shifts between implanted source current and recorded potential. However, no amplitude change would be expected, and in fact no amplitude change was observed. In various similar experiments, phase changes have been small or zero (Nunez and Srinivasan, 2006). To create amplitude reductions with an artificial dipole, one might employ adjustable pole separations in the artificial dipole. Similarly, the membrane scale capacitive-resistive effect outlined here involves a physiological reduction in all source-sink separations at higher frequencies, leading to a corresponding (low pass) reduction in **P**(**r**, *t*) and recorded EEG potential. This description is not accurate for ECoG, which is believed to record mainly local monopole sources; thus, we anticipate a reduced or perhaps zero low-pass columnar effect in cortical surface potentials.

### 5.3 Filtering by macro scale cortical asynchrony

This predicted low pass internal columnar effect (due to reduced source-sink separations) is quite distinct from the tangential influence of synchronous macro source regions **P**(**r**, *t*) over the cortical surface. Larger synchronous dipole sheets caused by neocortical dynamic interactions (shown in region a-i of fig. 4), may have minimal effects on ECoG magnitudes, but are expected to result in much larger EEG magnitudes. Such large changes in EEG amplitudes that occur when brain state changes are believed to be due mostly to large scale tangential synchrony changes; thus EEG scientists and clinicians have adopted the label *desynchronization* to indicate amplitude reductions, particularly in the case of alpha band rhythms, which seem to consist of mixtures of source regions of different sizes (Pfurtscheller and Lopes da Silva, 1999). “Synchrony” may refer to several distinct phenomena, occurring in different directions over the cortical surface and at different spatial scales. Source synchrony has been studied with both EEG data and volume conduction models, leading to several generalizations that appear to apply across a broad range data and head models (Nunez and Srinivasan, 2006). For fixed source strength **P**(**r**, *t*), we expect modest changes in scalp potential magnitudes as a dipole sheet, composed of synchronous macro sources **P**(**r**, *t*), grows from the sub-mm scale (diameter) to several mm. This is expected because the source region remains essentially a single dipole over this separation range from the distant locations of scalp measurements. As the dipole layer enlarges to diameters in the approximate range of about 1 to 10 cm, large increases in scalp potential magnitudes are expected as modeled in section 6. Scalp potential magnitudes may actually begin to decrease when dipole layer diameters become larger than about 20 cm due to the canceling effects of the curved head surface. Again, these arguments support the idea that *dynamic measures of cortical function are generally scale-dependent, and measures like functional connectivity cannot be interpreted in absolute terms*.

## 6. The forward problem: Scalp potentials due to macro sources

### 6.1 Sources confined to cortex

The macro source function or dipole moment per unit volume **P**(**r**, *t*) may be viewed as a continuous function of cortical location **r**, measured in and out of cortical folds as shown in fig. 4. **P**(**r**, *t*) generally forms a dipole sheet (dipole layer) covering the entire folded neocortical surface. Localized macro source activity is then just a special case of this general picture, occurring when **P**(**r**, *t*) is small or negligible at most cortical locations **r** due to any or all of the following properties of the micro sources *s*(**r**, **w**, *t*): (1) The micro sources are not sufficiently strong and/or not many are active. (2) The micro sources are asynchronous or randomly mixed across the depths of columns (coordinate **w**). However, even when **P**(**r**, *t*) is large in local cortex, it may not appear in the recorded EEG due to the volume conduction effects outlined below. Most scalp EEG is believed to be generated in neocortex; in such cases, the potential *V*(**x**, *t*) at scalp locations **x** may be expressed as double integral over the entire cortical surface area *A*,

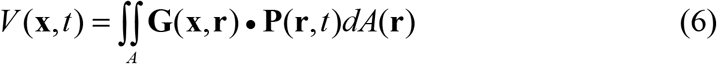

Here *dA*(**r**) is an element of cortical surface, and Eq (6) is a generalization of the basic dipole formula of Eq (2), but summed over many macro sources. The tissue boundaries and conductive properties of the head volume conductor are accounted for by the *surface Green’s function* **G**(**x**, **r**), which weighs the contribution of the macro source field **P**(**r**, *t*) according to macro source location **r** and the scalp recording point **x**(Jackson, 1975). When only a single isolated source occurs, **P**(**r,***t*) is a delta (point) function and **G**(**x**, **r**) is equal to the scalp potential due to that point source. The scalp potential due to many isolated sources or distributed sources is then just a sum (or integral) of all sources contributions, as given by Eq (6).

Because of the columnar structure of neocortex (Szentagothai 1978, 1987; Mountcastle, 1998; Nunez, 1995), we generally expect the vector **P**(**r**, *t*) to be pointed perpendicular to the local cortical surface (both folded and smooth), although Eq (6) does not require this condition. The surface integral in Eq (6) is a special case of the equivalent volume integral, the latter used when sub-cortical macro sources are included as possible inverse solutions (Nunez and Srinivasan, 2006). In the more general case, the surface elements *dA*(**r**) are replaced by voxels of mostly arbitrary size; the spatial scale of possible representative sources might range from several mm3 to 10s of cm3. However, as discussed in section 3, the dipole model is not valid for action potential sources in myelinated axons, which may account for the brainstem auditory evoked potential or other non-dipole phenomena (Nunez, 1995; Nunez and Srinivasan, 2006).

### 6.2 Effects of volume conduction

The Green’s (weighting) function **G**(**x**, **r**) will be small when the *electrical distance* between scalp location **x** and macro source location **r** is large. In an infinite, homogeneous, and isotropic (direction independent) medium the electrical distance equals the physical distance. But, due to volume conduction in the head, the two distances can differ substantially because of current paths distorted by variable tissue conductivities. Furthermore in anisotropic volume conductors, **G**(**x**, **r**) must also account for the fact that the conductivity of genuine tissue is direction dependent, resulting in a tensor or matrix form for **G**(**x**, **r**). While the accuracies of even the most sophisticated computer methods are limited by our limited knowledge of tissue conductivities and boundaries, approximate solutions of Eq (6) have proven to be quite useful in many EEG studies, mainly because many predicted effects are relatively insensitive to head model errors (Nunez and Srinivasan, 2006).

Contributions from different cortical regions may or may not be negligible in different clinical or cognitive studies. For example, source activity in large parts of mesial (underside) cortex and the longitudinal fissure (separating the brain hemispheres) may make negligible contributions to scalp potential in most brain states. Exceptions to this picture may occur in the case of mesial sources contributing to potentials at an ear or mastoid reference, an influence that has sometimes confounded clinical interpretations of EEG (Ebersole, 1997; Schomer and Lopes da Silva, 2018). In theory, a small number of synchronous macrocolumns could possibly produce sufficiently large dipole moments to be scalp-recorded. However, even in epileptic patients, it appears that **P**(**r**, *t*) rarely, if ever, reaches such magnitudes. The usual rule of thumb in clinical work with spontaneous EEG is that something like 6 to 10 cm^2^ of contiguous tissue (about 300 macrocolumns) must be synchronously active to be recorded as EEG (Cooper et. al, 1965; Delucchi, 1975; Ebersole, 1997; Nunez and Srinivasan, 2006; Schomer and Lopes da Silva, 2018). These estimates do not necessarily apply when scalp data are averaged over multiple trials in evoked (EP) or event related potentials (ERP).

### 6.3 Incorrect dipole assumptions

One may choose to define a single “dipole” (macro source) as the source activity in cm scale tissue volumes *W*; however, such action violates the dipole assumption required in the forward solution of Eq (6). In other words, *it appears that any EEG signal recorded on the scalp can be automatically excluded from the single (macrocolumn scale) dipole category unless its relative strength has been substantially enhanced by averaging over multiple stimulus trials.* Thus, we conclude that isolated “dipole sources” found as solutions based on non-averaged scalp potentials should be regarded as *representative* rather than *equivalent* or *genuine*. Such representative sources are perhaps best viewed as data reduction tools, somewhat divorced from the real physiology of neural sources. Apparently, all non-averaged EEG require extended dipole sheets of moderate to large sizes to be accepted as physiologically realistic. On the other hand, dipole localization algorithms that find the centers of extended dipole sheets can be useful, subject to proper interpretation. Another test of such algorithms is the predicted ratio of cortical to scalp potential. With a single macrocolumn scale dipole, this ratio should be large, perhaps something like 50 to 100 or more as suggested below in section 6.4. In contrast, experimental EEG/ECoG comparisons find ratios typically in the 2 to 5 range in spontaneous EEG (Cooper et al, 1965; Nunez and Srinivasan, 2006), again suggesting that recordable EEG sources typically form large dipole sheets.

### 6.4 Modeling scalp potential magnitudes with head models

These semi-quantitative arguments concerning the effects of synchronous dipole layer sizes on scalp potential magnitudes may be demonstrated with volume conductor models based on the general forward solution given by Eq (6). The standard head model employed here consists of four concentric spherical shells representing brain, CSF, skull, and scalp, typically labeled the “4-sphere” model (Nunez 1995; Nunez and Srinivasan, 2006). In our simulations, sphere radii are (brain, CSF, skull, scalp) = (8.0, 8.1, 8.6, 9.2 cm). Brain and scalp are assumed to have equal conductivities; CSF conductivity is assumed to be five times larger than brain conductivity. Brain-to-skull conductivity ratios in the range 20-160 are plotted here for historical reasons, but the lower range (20-40) is now considered more realistic. In any case, the main results reported here are relatively insensitive to model parameters. This is an important point since the 4-sphere model represents only a very approximate head model due mainly to our poor knowledge of tissue conductivities. This ignorance also places severe limitations on the more accurate geometric methods that employ finite element or boundary element models.

Figure 5 models scalp potential over the center of the spherical cap (located at the “north pole”) as a function of the sizes of cortical dipole layers (spherical caps). These dipole layers are formed by macro sources **P**(**r**, *t*) of constant strength in the 4-sphere head model (Nunez and Srinivasan, 2006). The idealized sources are assumed to be fully synchronous throughout the dipole layers and pointed perpendicular to the spherical cortical surface. Thus, sources in cortical folds are idealized as absent and/or canceling on opposite sides of fissures and sulci. Our source distributions consist of radial dipole layers located 1.4 cm below the outer scalp surface and forming spherical caps, regions of the spherical cortex which lie above a given plane. For all skull conductivity ratios, the predicted scalp potential is shown to be maximum for cap radii between about 5 and 15 cm (synchronous dipole layer diameters in the 10 to 30 cm range). Plotted scalp potential over the center of the spherical caps (north pole) is expressed as a percentage of estimated transcortical potential rather than **P**(**r**, *t*) because the former is often measured in cortical depth. While actual scalp magnitudes depend on scalp to skull conductivity ratios, the peak plot locations indicated in fig. 5 are insensitive to this parameter. In animal experiments employing depth recordings, transcortical potentials in the 200 μV range have often been reported, although such measures vary with experimental conditions (reviewed in Nunez, 1995). Thus, to the extent that these animal experiments pertain to human neocortex, fig. 5 suggests maximum human scalp potentials generated by large synchronous dipole layers in roughly the 100 μV range. In contrast, small (mm scale) dipole layers are expected to generate scalp potentials less than a few μV. The important influence of the reference electrode is ignored in these simulations, but the general ideas apply to most reference choices.

**Fig. 5.**
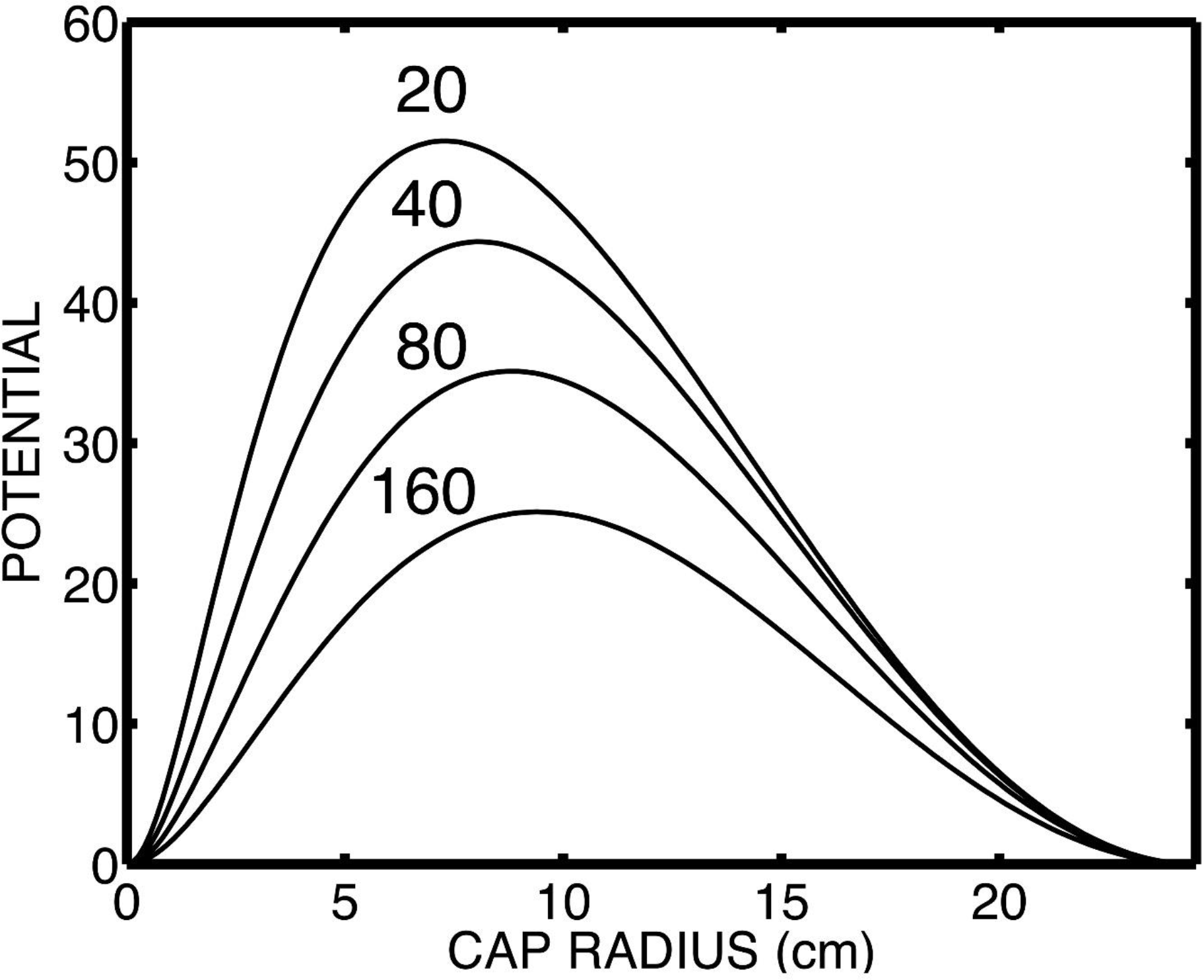
Normalized scalp potentials due to dipole layers of varying extent are plotted. The dipole layers form superficial spherical caps and are modeled with Eq (6) using a “4-sphere” model of the head. Each curve (vertical axis) shows predicted scalp potential over the center of the spherical cap (north pole), expressed as a percentage of (constant) transcortical potential. The horizontal axis is cap radius, the dipole layer size as measured from the “north pole” along the spherical scalp surface. The various plots indicate different brain to skull conductivity ratios. Most recent estimates of this effective ratio are in the 20-40 range.

## 7. The EEG inverse problem

### 7.1 Limitations of the inverse problem

The basic *inverse problem* in EEG is to employ measurements of potential distribution on the scalp surface *V*(**x**, t) to “invert” Eq (6); that is, to solve this integral equation for the macro scale function **P**(**r**, *t*) by employing a head model that provides the Green’s function **G**(**x**, **r**). Published descriptions of the inverse problem often emphasize different computer algorithms employed to produce approximate solutions based on different assumptions about brain physiology. But, none of these issues alter the physical fundamentals—a very large number of different source functions **P**(**r**, *t*) will yield the same surface potential distribution *V*(**x**, t) in any particular volume conductor. This is true whether or not the sources are assumed to be confined to neocortex as indicated by Eq (6). The source indeterminacy in EEG (or MEG) is fundamental; that is, it does not result from imperfect head models, noise, or under sampling, but occurs whenever the available data are limited to surface potentials. Given the non-uniqueness of the inverse problem, any inverse solution must depend partly on added information, the *solution constraints*. The origins of these constraints in published works have included plausible physiology-based conjectures, independent information obtained from MRI, fMRI or PET data, hopeful assumptions, or combinations of constraints (Nunez and Srinivasan, 2006); some of these are:

- The macro source function **P**(**r**, *t*) is assumed to be negligible at all but a few discrete locations. In other words, the sources are forced to consist of one or perhaps several isolated dipoles.
- All macro sources are assumed to be located in the crowns of cortical gyri, ignoring sources in cortical folds. Such solutions are expected to approximately match dura imaging or Laplacian estimates, discussed in section 8.
- Applying spatial smoothness criteria to the function **P**(**r**, *t*), or perhaps finding the inverse solution that optimizes (in some sense) the smoothness of **P**(**r**, *t*) (Pascual-Marqui, et al, 1994, 1999).
- Applying temporal smoothness criteria to the function **P**(**r**, *t*) (Scherg and von Cramon 1985). The possible justification is that sources are unlikely to turn on and off too abruptly.

### 7.2 Dipole localization

In practice, many spatial patterns of EEG can be approximately fit by a few dipoles. Other than in cases where the experimenter has independent evidence for isolated dipoles, perhaps early sensory evoked potentials, “equivalent dipoles” are more likely to be more accurately characterized as “representative dipoles.” For example, the most accurate dipole solutions may reflect centers of complex patterns of distributed cortical activity. Such representative information can be valuable, perhaps by informing epilepsy surgeons where to place the centers of cortical surface or depth arrays. More accurate source information may then be obtained with ECoG or iEEG.

Inverse EEG and MEG solutions involving single implanted dipoles in both physical head models and living human brains have successfully located these artificial sources within about 1-2 cm (Cuffin et al, 1991; Cohen et al, 1990; Leahy et al, 1998). However, when the sources are not known in advance to be exclusively isolated, inverse solutions face severe challenges because distributed cortical sources cannot normally be distinguished from isolated sources based only on scalp potential data (Nunez and Srinivasan, 2006). For this reason, many applications assume that all sources are cortical so that Eq (6) applies. This approach seeks distributed solutions by using thousands of dipole sources whose positions and orientations are fixed by the folded cortical surface. The problem is then to solve this under-constrained problem of determining the strengths of the dipole sources (Russell et al. 1998, 2005). But, this limited constraint does not, by itself, allow one to distinguish sources in cortical folds from source distributions in nearby cortical gyri that produce identical potentials on the dura and scalp surfaces. These two source categories are “equivalent” only in the limited sense of producing identical surface potentials.

This practical case of distinguishing sources in folds from gyri sources is easily explained. Any source distribution in a cortical fold will produce some potential distribution on the dura surface. In the absence of source constraints, one can easily find some extended source distribution in nearby cortical gyri that matches the dura potential due only to the deeper source region. For example, if we stick with assumed cortical sources normal to the dura surface, the resulting dura surface map will closely match the source map. Since there are no sources located between the dura and scalp surfaces, the scalp potential is then uniquely determined by Laplace’s equation given the dura potential (lower boundary condition). Additional information, perhaps obtained with MEG recordings, may allow the two kinds of source distributions to be distinguished. Another approach is to apply some spatial smoothness constraint to sources confined to the cortical tissue (Pascual-Marqui and Gonzalez-Andino, 1988; Pascual-Marqui et al, 1994; Pascual-Marqui, 1999); the resulting accuracy will, of course, depend on how well genuine sources conform to this constraint. Aside from uniqueness and source constraint issues, all inverse solution accuracies are limited by our imperfect knowledge of head volume conduction expressed by the Green’s function **G**(**x**, **r**) in Eq (6).

## 8. High resolution EEG

### 8.1 The Laplacian as a spatial filter

The label “high-resolution EEG” (HR EEG) refers to computer methods that spatially filter the scalp-recorded EEG, thereby effectively “reversing” some of the space-averaging caused by volume conduction between dura and scalp. The goal is to obtain more accurate estimates of the underlying dura potential rather than source localization. However, we argue here that high resolution EEG may also be viewed as a method of finding and characterizing *representative cortical sources*, which in cases of high electrode density, may be as close or closer to *equivalent sources* as the *representative sources* revealed by the inverse solutions of Eq (6). Our arguments are based on the underlying biophysics, not the specific computer algorithms employed. We confine the description to Laplacian methods, which are largely independent of head model, as opposed to model-dependent dura imaging. Our discussion is also confined to methods with moderate to high accuracy—spline Laplacians with at least 64 (preferably 128+) electrodes. While nearest-neighbor algorithms (Hjorth, 1975; Nunez, 1981, 2011; Nunez and Srinivasan, 2006) can be useful in a limited number of applications, our conclusions and recommendations apply only to splines obtained with dense electrode arrays. We also avoid the erroneous label “current source density (CSD)” in connection to HR EEG, which suggests a misleading connection to second derivative estimates in cortical depth recordings. In contrast to CSD studies, HR EEG (e.g. Laplacian) accuracy depends on the presence of high resistivity (low conductivity) skull layers that tend to focus currents perpendicular local skull tissue.

### 8.2 What does the Laplacian measure?

Laplacian methods provide reference-independent estimates of the dura surface potential based on scalp potential distribution. In contrast to inverse solutions, the relationship between dura and scalp potentials is unique, provided there are no sources located in between the two surfaces, and perfect knowledge of potential on an entire outer closed surface is obtained. While the latter idealistic condition does not hold in practice, the scalp surface Laplacian (*Lap*) has been shown to satisfy an approximate relationship to dura (or inner skull) potential that is entirely independent of reference electrode and mostly independent of head model (Nunez and Srinivasan, 2006). Let subscripts be as follows: (C, inner skull), (K, outer skull), and (S, outer scalp). The scalp Laplacian (*Lap*) is approximately proportional to the potential difference between outer and inner skull surfaces (*V*_*K*_ − *V*_*C*_):

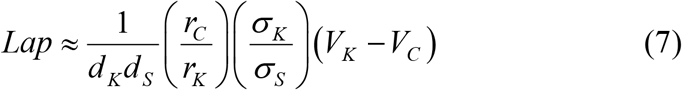

Here *d*_*K*_ and *d*_*S*_ are skull and scalp thicknesses, respectively, each roughly in the range 0.5 to 1 cm. *r*_*C*_ and *r*_*K*_ are radii of curvature of the inner and outer skull surfaces, the ratio is about 7/8, and is only weakly dependent on head shape. *σ*_*K*_ and *σ*_*S*_ are the conductivities of skull and scalp tissues; estimates of effective scalp to skull conductivity ratios are typically in the 20 to 80 range. The caveat “effective” accounts for corrections due to missing tissue layers or other approximations used in various head models. Such head models predict only small potential changes through the scalp thickness; thus, *V*_*K*_ is approximately equal to scalp surface potential. Models containing single dipoles also predict that the potential on the inner surface of the skull *V*_*C*_ is something like fifty to a hundred times larger than the potential on the outer surface of the skull *V*_*K*_. Thus, in the case of single dipoles (or small dipole layers), the magnitude of the outer skull surface potential*V*_*K*_ is negligible in comparison to the inner skull surface potential *V*_*C*_. It then follows from Eq (7) that, in the case of localized cortical source regions, the (negative) surface Laplacian (*Lap*) is proportional to the potential on the inner surface of the skull *V*_*C*_. In cases of a thin CSF, dura potential is approximately equal to *V*_*C*_.

Whereas this analysis is based mainly on physical arguments and computer simulations (Nunez, 1981, 2011; Nunez et al, 1994; Nunez and Srinivasan, 2006), its moderate to high accuracy in predicting dura potential distributions has been confirmed in more detailed mathematical and simulation studies (Perrin et al, 1987, 1988; Pascual-Marqui et al, 1988; Nunez et al, 1991; Law et al, 1993; Babiloni et al, 1996; Kramer and Szeri, 2004; Nunez and Srinivasan, 2006; Kayser and Tenke, 2015). Furthermore, the realistic scalp geometry obtained from MRI has been incorporated in surface Laplacian estimates that take into account local scalp curvature (Deng et al, 2012). Our reference to Laplacian “accuracy” indicates only that the proportionality indicated in Eq (7) holds approximately; however, no claim is made that the actual dura potential measured in μV can be estimated accurately. The effective scalp to skull conductivity ratio (*σ*_*S*_ / *σ*_*K*_) is poorly known; it probably varies over the scalp surface of individual subjects, across subjects, and more. However, in most, if not all, applications we are most interested in the cortical locations of Laplacian peaks and not especially interested in their actual peak magnitudes. This same limitation concerning absolute strengths applies to all inverse solutions and high resolution methods; predicted source magnitudes are sensitive to the scalp to skull conductivity ratio, which may be uncertain by a factor of three or more (Nunez and Srinivasan, 2006).

### 8.3 Simulated Laplacians and cortical potentials

A simple Laplacian simulation, based on the 4-sphere head model, is shown in fig. 6 (Nunez and Srinivasan, 2006). Two radial dipoles **P**(**r**, t) are located at a depth of 1.4 cm below the scalp surface, corresponding to sources in cortical gyri. One radial dipole is oriented with the positive pole up (solid contour lines) and the other radial dipole has the negative pole up (dashed contour lines). The third dipole is tangential, as indicated by the positive (+) and negative (-) poles, and located at a depth of 2.2 cm below the scalp, simulating an isolated dipole in a sulcal wall. The three examples (a-c) correspond to increasing tangential dipole strengths—(a) the three strengths are equal, (b) the tangential dipole is twice as strong as the radial dipoles, (c) the tangential dipole is four times as strong. In case (a), the two radial dipoles provide a much stronger contribution to surface potential than the tangential dipole, with only a weak positive region evident over the right side of the map. The potential generated by the negative radial dipole appears to fall off with tangential distance more slowly than the positive dipole and have larger magnitude due to the influence of the tangential dipole.

**Figure 6.**
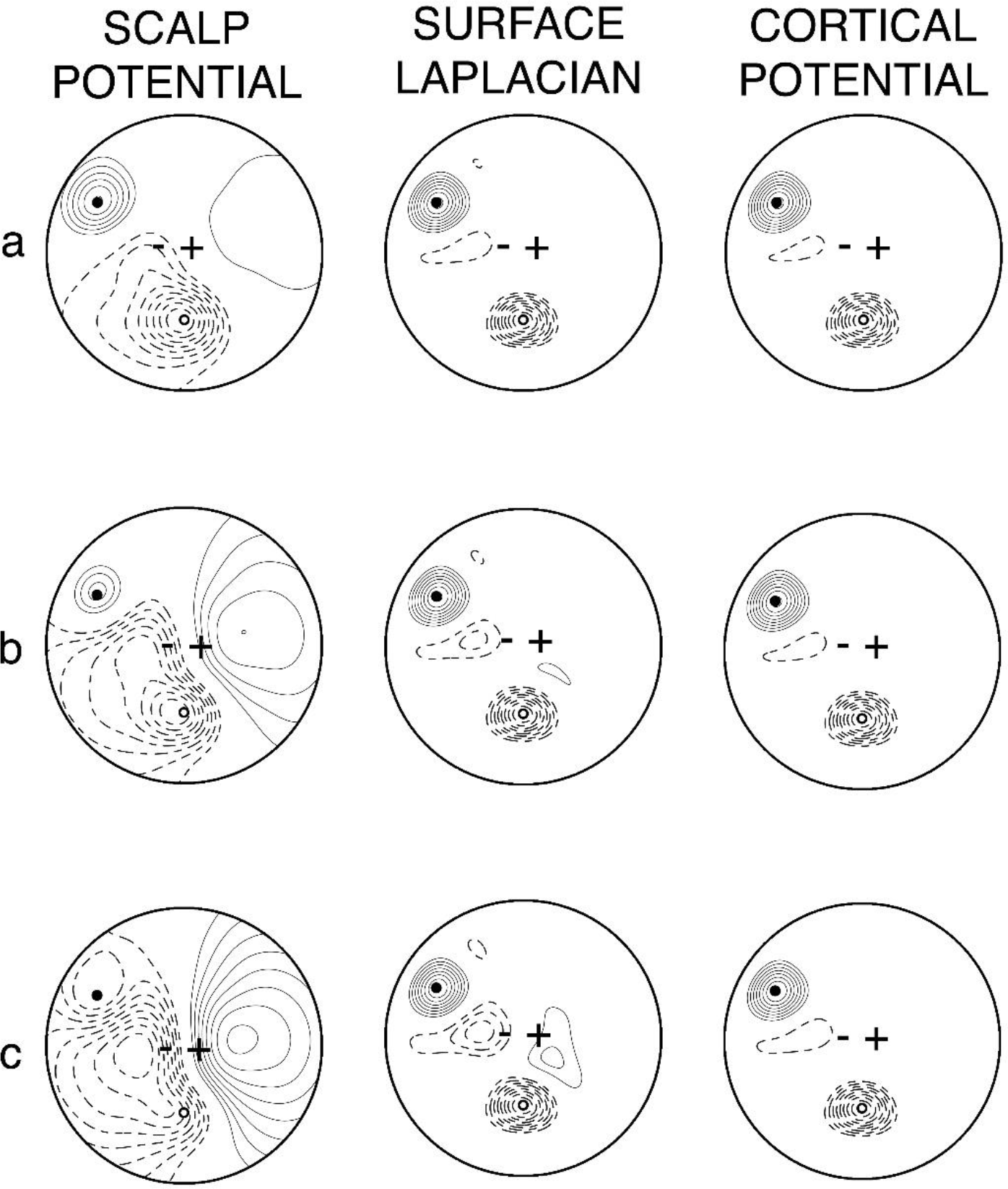
Simulated scalp potential, Laplacian, and cortical potential due to three dipoles **P**(**r**, *t*), two radial, one tangential, in a four sphere head model. The ratios of tangential to radial dipole strengths are (a) one, (b) two, (c) four. Reproduced with permission from Nunez and Srinivasan, 2006.

The surface Laplacian and cortical potential maps are nearly identical as predicted by Eq (7), revealing positive and negative radial dipoles with nearly equal magnitudes, but these maps do not reveal the tangential dipole. In case (b), the broad field of the stronger tangential dipole becomes evident on the right side of the potential map, while a complex distribution appears over the left side, reflecting a mixture of contributions from both radial and tangential dipoles. The surface Laplacian and cortical potential again identify the two radial sources with magnitude unaffected by the tangential dipole. In case (c), the potential distribution is dominated by the single tangential dipole. The Laplacian again reveals the two radial dipoles, but also detects a much smaller field associated with the tangential dipole.

The effect of dipole layer size on sensitivity to the Laplacian is also estimated with the 4-sphere head model as indicated in fig. 7. The dipole layers are identical to those shown in fig. 5, which indicates the sensitivity of unprocessed scalp potential to dipole layer size. The implications of these plots may be demonstrated with a thought experiment in which cortical synchrony slowly spreads uniformly in all directions from some isolated cortical region, e.g., a mm size patch at the north pole of the model sphere. The small isolated cortical patch produces a scalp potential that is probably too small to measure, at least without averaging. As the region of synchronous sources (the dipole layer) expands to something like the 1 cm scale, the Laplacian over the center of the layer grows rapidly, whereas the unprocessed potential is likely to remain masked by distant sources and artifact. As indicated in fig. 7, as the diameter of the region of synchrony expands beyond the 10 cm scale (cap radius of 5 cm), the Laplacian magnitude falls off sharply, whereas the unprocessed potential (fig. 5) continues to increase with spreading synchrony until the dipole layer diameter expands to approximately the 15 cm range.

**Fig. 7.**
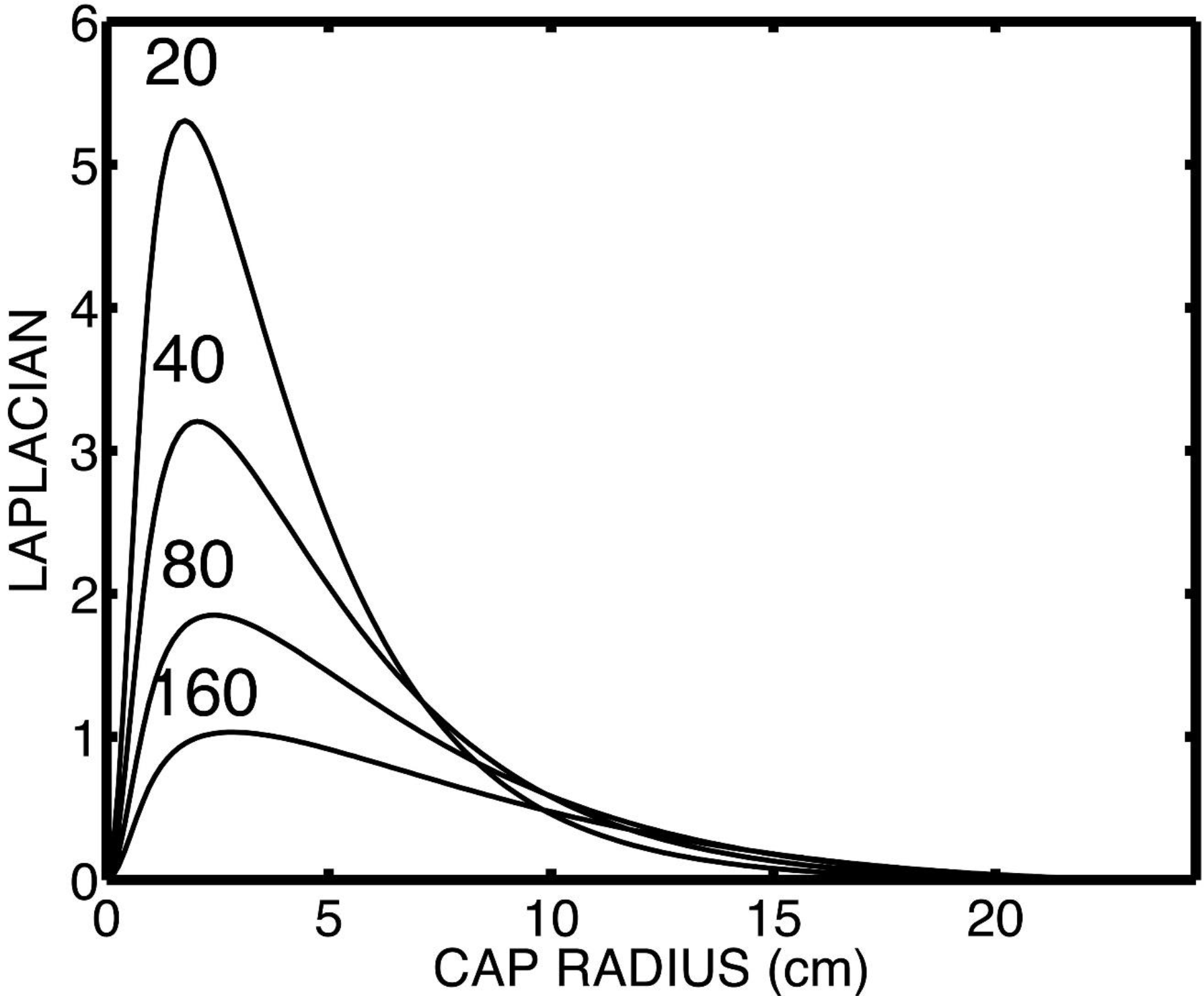
The scalp Laplacian is modeled as a function of dipole layer size. The same spherical caps used in fig. 5 are employed here. The Laplacian (vertical axis) is expressed as a percentage of transcortical potential.

The simulations of figs. 5-7 indicate that the unprocessed potential and Laplacian are selectively sensitive to synchronous cortical regions of different sizes; they provide complementary, but partly overlapping measures of cortical dynamics. In particular, the effectiveness of the surface Laplacian as a tool to identify EEG sources depends strongly on the spatial properties (depth and orientation) of the sources. *If localized and superficial radial sources are not the primary contributors to any particular EEG signal, the surface Laplacian will tend to filter out much of the signal.* One possible example of such large scale synchrony is the classic spike and wave occurring in epileptic patients undergoing petit mal seizures (Sharbrough and Lagerlund, 1990).

### 8.4 Dura imaging, comparison with Laplacians

Dura imaging algorithms provide inner continuation solutions. In contrast to the non-unique inverse problem, dura imaging is based on the unique relationship between potentials on any two closed surfaces in a volume conductor when no sources are located in between the surfaces (Cadusch et al, 1992; Nunez et al, 1994). How do such dura image results compare with Laplacian estimates of dura potential maps? The answer depends on electrode density and head model accuracy. Laplacians are mostly independent of head model, but rely on dense electrode arrays. Accurate head models, if actually achieved, could possibly allow dura imaging to provide accurate estimates of dura potential with lower electrode density. We are aware of only one set of studies directly comparing the two HR EEG methods (Nunez et al, 2001; Wingeier, 2004; Nunez and Srinivasan, 2006, chapter 8). With 64 electrodes, correlations between the New Orleans spline Laplacian and Melbourne dura imaging were found to be in the 0.8 range *in both recorded EEG and simulations*. In similar studies with 131 electrodes, correlations were in the 0.95 range in *both recorded EEG and simulations*. The EEG was standard eyes closed alpha rhythm. The simulations involved 100 different source patterns, each consisting of 3602 cortical dipoles **P**(**r**, *t*) in a three-concentric spheres head model. *The two HR EEG methods agreed even though dura imaging was based on average reference recordings, whereas the Laplacian was entirely reference free.*

In summary, the above cited correspondence between Laplacian and dura potential is not valid if the cortical sources are too broadly distributed because in such cases *V*_*K*_ and *V*_*C*_ are in the same general range. For very large dipole sheets, *V*_*C*_ may be only 2 to 5 times as large as *V*_*K*_, a ratio range consistent with much spontaneous EEG data (Cooper, et al, 1965; Nunez and Srinivasan, 2006). In summary, the surface Laplacian provides plausible semi-quantitative estimates of dura surface potential when the underlying source regions are relatively localized, with diameters smaller than perhaps 10 cm. By contrast, very broadly distributed source distributions (dipole layers) cause the surface Laplacian to underestimate dura surface potentials. Expressed another way, the Laplacian is relatively insensitive to very low spatial frequency source activity. It tends to filter out such sources, along with low spatial frequency potentials due only to volume conduction.

### 8.5 Source localization versus Laplacian estimates

This selective sensitivity of the surface Laplacian to localized source regions appears to be of substantial importance when the main objective is source localization, as in the example of focal epilepsy. The Laplacian is independent of reference electrode choice or a priori assumptions about sources. It is also largely independent of volume conductor model, other than the assumption of low conductivity skull layers. In practice, the accuracy of Laplacian estimates is limited by noise, under sampling, and inhomogeneous skull properties; however, most indications suggest that it reliably distinguishes isolated cortical sources from broadly distributed source patterns. Thus, we suggest that when high density electrode arrays are employed (64 to 128+), attempts at source localization based on recorded scalp potential distribution could begin productively with a spline Laplacian estimate. In the general use of EEG data in cognitive and clinical studies, the surface Laplacian provides a spatial filtering of the EEG that limits electrode sensitivity to “local” sources (within a few cm of each electrode), thereby revealing source dynamics at smaller spatial scales than scalp potentials (Nunez, 1995; Nunez et al, 1997, 1999; Nunez and Srinivasan, 2006; Srinivasan, 1999; Srinivasan et al, 2007). *However, if the surface potential patterns of interest (perhaps broad peaks of potential) disappear when the Laplacian is applied, this outcome provides strong evidence that the potential patterns are generated by widely distributed and/or non-superficial sources (e.g., in cortical folds), not localized gyri sources.* Any algorithm that “finds” local superficial cortical sources in such a case would be suspect; such solutions should be viewed as *representative*, not *genuine* or even *equivalent*.

## 9. Conclusions

### 9.1 Source interpretations of brain patterns

Brains exhibit multiscale dynamic source patterns, essentially the electrical *signatures* of mental states. Source information may include ERP magnitude changes and latencies from sensory stimuli or changes in EEG oscillation frequencies in extended networks. Signatures also include various kinds of correlations between different parts of the brain—coherence, covariance, Granger causality, functional connectivity, and so forth (Sporns, 2011). Any measured correlation between a pair of signals is likely to be sensitive to the chosen correlation measure. For example, two signals can exhibit quite different coherence in different frequency bands as demonstrated by alpha rhythm and other data (Nunez, 1995; Nunez et al, 1997, 1999, 2015; Srinivasan et al, 2007). Or, a pair of ERP signals recorded at different locations and associated with some task can show quite a different covariance depending on lag time between scalp locations (Gevins et al. 1994, 1997). This general outcome emphasizes that source networks can act locally in some ways, but at the same time, act globally in other ways, thereby directly addressing the *binding problem* of brain science (Nunez, 1989, 2012, 2016; Nunez and Srinivasan, 2006, 2014). In other words, signals and their source networks may be “functionally connected” in some ways, but at the same time, “functionally disconnected” by other measures. In essence, different parts of the brain do different things, but these sub systems also act together in healthy brains to produce an integrated consciousness (Silberstein, 1995; Edelman and Tononi, 2000).

### 9.2 Micro and macro sources

The conceptual framework proposed here is aimed at electrophysiological studies of brain source activity recorded at distinct tissue scales, including LEP and ECoG, but with emphasis on EEG. Our purpose is to facilitate proper interpretations of recorded potentials in terms of the underlying sources. At smaller scales, electric potentials are generated by micro scale current sources *s*(**r**, **w**, *t*). For large scale EEG studies, macro sources **P**(**r**, *t*) are defined for tissue volumes containing many micro sources *s*(**r**, **w**, *t*). When defined at the mm scale of cortical macrocolumns, **P**(**r**, *t*) is essentially the “effective” (diffuse) current density through the local cortex. **P**(**r**, *t*) may be treated as a continuous field variable that generally forms a dipole layer (sheet) over the entire (folded) cortex. Localized cortical activity is then just a special case, occurring when **P**(**r**, *t*) is negligible or small at most locations. This theoretical treatment of tissue current sources is closely analogous to the classical electromagnetic theory of physical materials (dielectrics) involving both micro and macro scale charge sources and fields.

The inverse problem is non-unique—many **P**(**r**, *t*) can produce the same surface potential distribution. Thus, in practice, inverse solutions must involve constraints; that is, assumptions or additional information imposed on solutions. One may constrain solutions to cortex, but this limited constraint does not, by itself, allow one to distinguish sources in cortical folds from source distributions in nearby cortical gyri that may produce identical potentials on the dura and scalp surfaces. These two source categories are “equivalent” only in the limited sense of producing identical surface potentials. With such issues in mind, we emphasized distinctions between *genuine sources*, *equivalent sources*, and *representative sources*. In practice, many spatial patterns of EEG can be approximately fit by a few “dipoles,” meaning the macro sources **P**(**r**, *t*). Other than in cases where independent evidence for isolated sources is known, so-called *equivalent dipoles* are more accurately characterized as *representative dipoles*. For example, even the most accurate inverse dipole solutions may reflect centers of complex patterns of distributed cortical activity.

### 9.3 Experimental and computational strategies

What are the “best” methods to record, analyze, and interpret EEG data in terms of the underlying brain sources? All methods are subject to issues mostly avoided here: artifact, reference, electrode density, head model, algorithm, and noise errors, but here is short list of several possible experimental strategies:

- <u>The classical approach</u>. Spatial transformation is limited to re-referencing, often the common average reference (Nunez and Srinivasan, 2006; Nunez, 2010a). In the past, some have viewed this approach as locating source regions, with one so-called “source” under each electrode, but such *representative sources* are generally not even *equivalent*, much less *genuine*.
- <u>Constrained inverse solutions estimating **P**(**r**, *t*)</u>. Accuracy is limited by the chosen constraints and head model. Simulations employing localized and distributed sources over a range of head models can provide some idea of when the methods work and when they break down. In such simulations, the test head model should differ in varying degrees from the head model used to obtain the inverse solution.
- <u>High resolution EEG, based on Laplacians or dura imaging</u>. The Laplacian enjoys the major advantages of being entirely independent of reference electrode and mostly independent of head model, but moderate to high accuracy requires dense electrode arrays (64-128+). Dura imaging relies on a head model, but might require fewer electrodes. The resulting dura potential estimates can be viewed as *representative* gyri sources, which can be *equivalent* only when all other sources, including those in cortical folds, can be neglected.
- <u>Combined dura imaging and Laplacian methods</u>. Both methods provide estimates of dura potential patterns, but are mostly independent of each other. Their magnitudes cannot be compared directly since they have different units; however, normalized comparisons indicate close agreement with both actual and simulated EEG when dense electrode arrays are employed (Wingeier, 2004; Nunez and Srinivasan, 2006).
- <u>Combined classical and Laplacian approaches</u>. The surface Laplacian provides spatial band pass filtering of EEG that limits electrode sensitivity to “local” sources, thereby revealing source dynamics at smaller spatial scales than raw scalp potentials. Maximum scalp potentials occur with synchronous dipole layers having diameters roughly in the 10 to 20 cm range, whereas Laplacians are most sensitive to dipole layers approximately in the 1 to 6 cm range as indicated in figs. 5 and 7 (Nunez and Srinivasan, 2006). By analyzing both measures in the same data sets, one can obtain complementary information about neocortical dynamics at different scales (Nunez, 1995; Nunez et al, 2015). In this manner, estimates of distinct multi-scale coherence patterns can be obtained (Srinivasan et al, 2007; Nunez et al 1997, 1999; Nunez and Srinivasan, 2006). Multi-scale coherence differences between adults and children provide just one example (Srinivasan et al, 1999).
- <u>Combined inverse solutions and Laplacian approaches</u>. Attempts at source localization may benefit from prior Laplacian estimates. If the surface potential patterns of interest (perhaps broad peaks of potential) disappear when the Laplacian is applied, such outcome provides evidence that the potential patterns are generated by non-superficial and/or widely distributed sources, not localized gyri sources. Any algorithm that “finds” local cortical sources in gyri in such case would be highly suspect; such inverse solutions are likely to be *representative*, not *genuine* or even *equivalent*.

## Acknowledgements

The authors would like to thank members of the IFCN Workshop, and especially its leader, Claudio Babiloni for extended and useful discussions of EEG issues. This research was supported by National Institutes of Health of the United States grant 2R01MH68004.

